# Spotted hyena gut cross-talks: Symbionts modulate mucosal immunity

**DOI:** 10.1101/2024.07.25.605121

**Authors:** Susana P. V. Soares, Victor Hugo Jarquín-Díaz, Miguel M. Veiga, Stephan Karl, Gábor Á. Czirják, Alexandra Weyrich, Sonja Metzger, Marion L. East, Heribert Hofer, Emanuel Heitlinger, Sarah Benhaiem, Susana C. M. Ferreira

**Author notes:** Max-Delbrück-Center for Molecular Medicine in the Helmholtz Association (MDC). Robert-Rössle-Str. 10, 13125 Berlin, Germany.

## Abstract

The intestinal mucosa is at the front line of host-microbiome interactions, but little is known about these interactions within natural populations. Here, we non-invasively investigated associations between the gut microbiome and mucosal immune measures while controlling for host, social, and ecological factors in 199 samples of 158 wild spotted hyenas (*Crocuta crocuta*) in the Serengeti National Park, Tanzania. We profiled the microbiome composition, including bacteria, fungi and parasites, using a multi-amplicon approach, and measured faecal immunoglobulin A and mucin. Probabilistic models indicated that both immune measures predict microbiome similarity among individuals in an age-dependent manner. The strength of the association effect varied, being strongest within bacteria, intermediate within parasites, and weakest within fungi communities. Machine learning regression accurately predicted both measures and identified the taxa driving these associations: symbiotic bacteria reported in humans and laboratory mice, unclassified bacteria, a hookworm, host DNA likely reflecting inflammation, and diet. Our findings indicate a complex interplay between the host, its environment and symbionts. These findings increase our knowledge of the gut microbiome in natural populations, which harbour highly dynamic and diverse eukaryotes under the influence of unpredictable environmental factors and where selection is not artificially biased.

## Introduction

The gut microbiome, the community of micro-and macroorganisms and their products or genetic material, residing within the gastrointestinal tract influences host physiology and plays a vital role in maintaining homeostasis and health ^1^. In turn, the host’s physiology, particularly the mucosal immune system, shapes this community. The intestinal mucosa, a single layer of epithelial cells covered by the mucus layer, is the first line of defence against pathogen translocation and fosters the maintenance of beneficial taxa ^2,3^. It produces and releases a variety of enzymes and various immune defence molecules, e.g. antimicrobial peptides, mucins, and antibodies, to maintain intestinal homeostasis ^4–6^. Perturbations of this community are associated with intestinal diseases, often accompanied by increased mucosal permeability and disrupted immune responses ^7–9^.

Two important and broad-acting measures of mucosal immunity are secretory immunoglobulin A (IgA) and mucins. IgA is the primary antibody secreted into the intestinal lumen. It coats the surface of a broad but defined subset of gastrointestinal taxa ^10^ and is a substantial metabolic substrate for microorganisms ^11,3^. IgA also prevents pathogens from crossing the intestinal epithelium and neutralises toxins and virulence factors ^12^. All these processes selectively promote or hamper the colonisation and growth of specific taxa, thereby supporting the intestinal barrier in the control of the gut microbiome. Mucins form a mucus layer that covers the epithelium, serves as a lubricant to protect against mechanical stress, and limits its direct contact with toxins, digestive enzymes and potential pathogens ^13,14^. Mucins are also important metabolic substrates for the gut microbiome, providing attachment sites that promote colonisation and provide a selective environment for symbionts ^15,14^. Despite a growing body of knowledge of interactions between intestinal mucosal immunity and microbiome, the focus remains disproportionately on bacteria ^16,17^.

Although bacteria outnumber eukaryotes within mammalian intestines ^18^, eukaryotes such as fungi, helminths and protozoa also play a significant role in regulating the gut microbiome ^19^ and host health ^20–22^. Studies on immune responses to fungi and parasites (i.e. protozoans and helminths) are often performed in isolation from the rest of the intestinal community, although inter-species interactions likely shape both the gut microbiome and immune responses ^23–25^. Host-microbiome interactions via mucosal immunity are poorly understood, and the knowledge gap is even wider in wild animal populations, where the acquisition of samples can be challenging and species-specific reagents and appropriate immune assay validations are limited ^26^.

Wild animals living in natural environments have a distinct microbiome compared to their captive counterparts ^27^, characterized by a higher diversity of eukaryotes ^28,29^. The study of host-microbiome interactions in wild animals has provided important insights into how host characteristics and the ecological environment shape their intestinal communities. Microbiome composition changes with age and between life stages, particularly during ontogeny, as observed in several mammals ^30–32^. This effect is often attributed to changes in host physiology, particularly immune development, but also behavioural and dietary changes. Additionally, the host environment (biotic and abiotic factors) shapes the composition of gut microbiomes, as seen by spatial and temporal heterogeneities ^33–35^, the effect of diet ^36,37^ and social interactions ^38,39^. Importantly, the environment interacts with host physiology and genetics, and these interactions might change over time and are as such context dependent ^40^. Long-term individual-based field studies allow the collection of detailed data on the life histories of individually recognised animals and on the biotic and abiotic environment, often with minimal anthropogenic manipulation. These are particularly suitable to disentangle the relative contributions of host characteristics and ecological factors in shaping microbiome composition and potential interactions within this community and with the host ^41–43^.

Here, we study a wild population of spotted hyenas (*Crocuta crocuta*) in the Serengeti National Park, monitored within a long-term research project, to investigate the links between intestinal mucosal immunity and gut microbiome. Spotted hyenas (hereafter ‘hyenas’) live in stable social groups (clans) with a linear dominance hierarchy (i.e. with social ranks) in which females and their offspring socially dominate immigrant males ^44,45^. Because they both hunt and scavenge and thus can consume fresh and highly decomposed carcasses ^46^, hyenas can be exposed to a variety of pathogens. Additionally, pathogen transmission among clan members is facilitated due to their high rate of social interactions ^47–49^. Nonetheless hyenas are known to be able to remain healthy ^47,50^, suggesting specific adaptations such as a specialised immune system and/or resilient intestinal community ^48,51^. Previous research has shown that host characteristics (e.g. age) and environment (e.g. clan membership) affect parasite infections ^49,52,53^ and intestinal bacterial composition ^30,54,55^. Furthermore, faecal IgA and mucin reflect *Ancylostoma* load, a parasite shown to reduce longevity in the Serengeti population ^49,56^. We hypothesised that the host-associated taxa are under tight regulation by mucosal immunity and are thus strongly associated with faecal IgA and mucin levels.

## Materials and methods

### Study site, population under study and sample collection

We collected life-history, behavioural and ecological data, and 210 faecal samples from 165 individually recognised wild spotted hyenas from 2004 to 2018, Figure 1A, Figure S1. Hyenas from this study belong to three clans monitored on a near daily basis since 1987, 1989, and 1990 in the context of an ongoing long-term individual-based research project in the Serengeti National Park (SNP), Tanzania. Individuals are recognised based on their unique spot patterns and other features, such as scars and ear notches ^44,57^.

**Figure 1.**
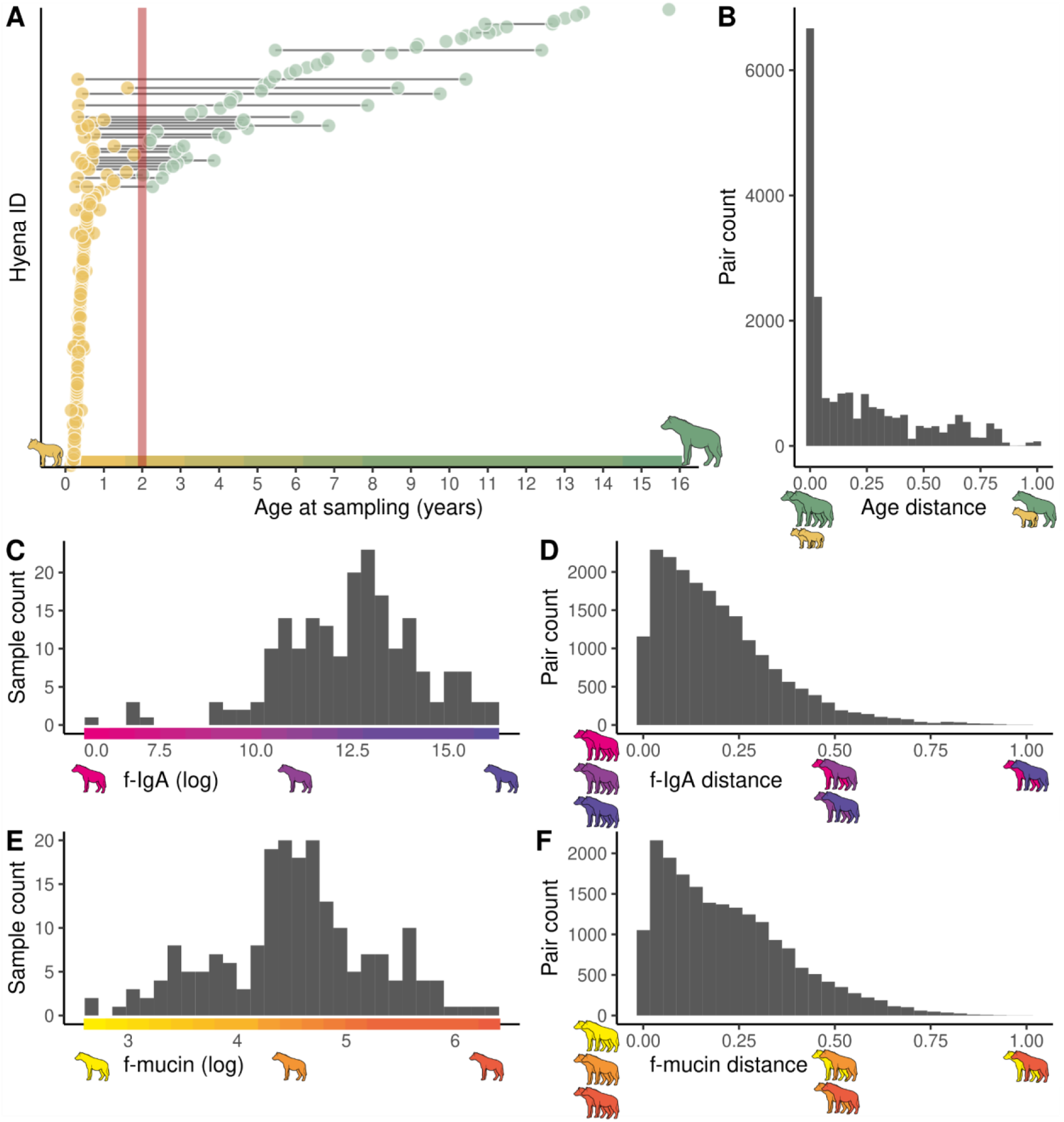
Distributions of age and mucosal immune parameters in 199 samples from 158 free-ranging hyenas used in this study. **A**) Age at sampling. Samples from the same individual are connected with a grey line and dots are coloured based on the age category (yellow: juveniles, green: adults). The red vertical line indicates the age threshold between categories (2 years). **B**) Distribution of the age distance of compared samples. **C**) Distribution of faecal IgA. **D**) Distribution of faecal mucin. **E**) Distances between faecal IgA concentrations of all compared samples. **F**) Distances between faecal mucin concentrations of all compared samples. RU: relative units. Hyena icons are colour coded to illustrate the ranges of age and immune measures levels in the sampled individuals.

Cubs were aged to an accuracy of ±7 days based on their behaviour, movement coordination, size and pelage when seen for the first time ^58^. By the age of approximately three months, sex was determined by the shape of the external genitalia, particularly the dimorphic glans morphology of the erect phallus ^59^. Maternal identity was determined based on nursing observations at the communal den(s) and was further confirmed by DNA microsatellite loci analysis ^60^.

The social rank of adult females in their clans was determined based on the observation of submissive acts in dyadic interactions recorded *ad libitum* and during focal observations ^61^. For each clan, we used the outcome of these dyadic interactions to construct an adult female linear dominance hierarchy that was updated daily after demographic changes (recruitment or deaths of adult females) and socially mediated changes in rank (coups). To make rank positions comparable across clans and within clans when the number of females in the hierarchy changed, we assigned standardised ranks, evenly distributing social ranks from the highest (standardised rank: +1) to the lowest rank (standardised rank: −1) within a clan. Juveniles (individuals younger than two years) were assigned the same standardised ranks as the mothers raising them at the sampling date ^62^. Six hyenas sampled before reaching adulthood were adopted or jointly raised by their genetic and surrogate mothers ^62^. In these cases, we assigned the social rank of the surrogate mother or the average of the rank of the mothers in the case of joint-raising.

Faecal samples were immediately collected after defecation and refrigerated in cool packs in the field until transport to the field station where they were mechanically mixed and aliquots stored at -10°C until transport to storage at - 80°C in Germany ^52^. Aliquots for DNA extraction were stored in RNAlater (Sigma-Aldrich, St Louis, MO, USA).

### Faecal immunological assays

Faeces aliquots were lyophilised for 22 hours in a freeze-dryer (Epsilon1-4 LSCplus, Martin Christ GmbH, Osterode, Germany) followed by homogenisation with mortar and pestles. Faecal mucin (f-mucin) and faecal immunoglobulin A (f-IgA) assays, previously adapted and validated for application to spotted hyena faeces, were applied using the methods described in detail in ^56^ and details are found in the Supplements.

### DNA extraction

We extracted genomic DNA from faeces using the NucleoSpin®Soil kit (Macherey-Nagel GmbH & Co. KG, Düren, Germany) under the manufacturer’s protocol with the following modifications: we performed mechanical lysis of the sample in a Precellys®24 high-speed benchtop homogeniser (Bertin Technologies, Aix-en-Provence, France) with two disruption cycles at 6000 rpm for 30 s, with a 15 s delay between them. We eluted DNA in a 40 µL TE buffer. Quality and integrity of the DNA were later evaluated with a full-spectrum spectrophotometer (NanoDrop 2000c; Thermo Fisher Scientific, Waltham, MA USA). We used a Qubit® Fluorometer and the dsDNA BR (broad-range) Assay Kit (Thermo Fisher Scientific) to quantify the concentrations of double-stranded DNA. The DNA extracts were adjusted to a final concentration of 50 ng/µl with nuclease-free water (Carl-Roth GmbH + Co. KG), and stored at −80°C, until further use.

### Library preparation and sequencing

We used faecal DNA preparations for multimarker amplification using the microfluidics PCR system Fluidigm Access Array 48 x 48 (Fluidigm, San Francisco, California, USA). We randomised sample order and amplified samples in parallel with non-template negative controls using a microfluidics PCR. This allowed the amplification of multiple fragments (amplicons) for different marker genes (primer pairs in additional supplementary file 1). We integrated PCR setup library preparation into the amplification procedure according to the protocol for Access Array Barcode Library for Illumina Sequencers (single direction indexing) as described by the manufacturer (Fluidigm, San Francisco, California, USA). The amplicons were quantified using the Qubit fluorometric quantification dsDNA High Sensitivity Kit (Thermo Fisher Scientific, Walham, USA) and pooled in equimolar concentrations. The final library was purified using Agencourt AMPure XP Reagent beads (Beckman Coulter Life Sciences, Krefeld, Germany). The quality and integrity of the library were confirmed using the Agilent 2200 TapeStation with D1000 ScreenTapes (Agilent Technologies, Santa Clara, California, USA). Sequences were generated at the Berlin Center for Genomics in Biodiversity Research (BeGenDiv) on the Illumina MiSeq platform (Illumina, San Diego, California, USA) using v2 chemistry with 500 cycles. All sequences are accessible in BioProject PRJNA1134446 in the NCBI Short Read Archive (SRA).

### Identification and quality screening of the amplicon sequence variants (ASVs)

All analyses were performed using R v 4.4.0 ^63^. For the initial analysis we used the packages dada2 v. 4.3.1 ^64^ and MultiAmplicon v. 0.1.1 ^65^ to filter, sort, merge, denoise and remove chimaeras for each run separately and for each amplicon, and removed contaminants with decontam v. 1.21.0 ^66^ using “prevalence” and “frequency” methods (method = ”combined”). We further removed amplicon sequence variants (ASVs) with less than 1% prevalence, less than 0.005% relative abundance ^67^ and samples with less than 100 reads. Each amplicon in the multi-amplicon datasets was individually filtered and then all products were collated into an “phyloseq” object with the function “merge_phyloseq” from the package phyloseq v. 1.45.0 ^68^. This last step resulted in 199 samples.

### Taxonomic annotation of ASVs

We used the RDP classifier ^69^ implemented within dada2 v. 1.29.0 package ^64^ to assign taxonomy to the resulting ASVs. Sequences targeting the 18S, 16S, 28S and ITS rRNA genes were classified against the SILVA 138.1 SSU Ref NR 99, the SILVA 138.1 LSU Ref NR 99 ^70^, the UNITE ^71^ databases, respectively. We used the SILVA 138.1 SSU Ref NR 99 to classify 18S and 16S rRNA gene sequences, the SILVA 138.1 LSU Ref NR 99 databases ^72^ for 28S rRNA gene sequences, and the UNITE database ^73^ for ITS rRNA gene sequences. All other sequences from targeted regions without publicly available curated databases were classified against sequences downloaded from NCBI using RESCRIPt ^74^.

### Merging ASVs into combined ASV (cASV)

Our final dataset had ASVs from different amplicons targeting different marker loci of the same taxon. Hence, we merged the ASVs originating from the same taxon on the basis of their co-abundance within each genus (n = 476). Co-abundance networks were constructed based on positive (Pearson coefficient > 0) and significant correlations (p < 0.01), after correction for multiple testing with the Benjamini-Hochberg method. ASVs that clustered together using the “cluster_fast_greedy” function from the igraph package ^75^ were then merged into one ASV by summing their abundances into combined ASVs (cASV). This has been previously accessed for *Eimeria* spp. ^76^ and is now extended to all annotated genera.

### Statistical analysis

We conducted all analyses using R 4.4.0 ^63^. The intestinal community composition was decomposed into four groups of cASVs: 1) bacteria domain, including the phyla Firmicutes, Bacteroidota, Campylobacterota, Cyanobacteria, Proteobacteria, Planctomycetota, Fusobacteriota, Actinobacteriota, Deferribacterota, Spirochaetota, Desulfobacterota and unclassified (unknown) bacteria; 2) fungi, including the phyla Mucoromycota, Ascomycota, Basidiomycota, Blastocladiomycota, Chytridiomycota, Neocallimastigomycota; and 3) parasites, including the known protozoa and helminth genera *Sarcocystis*, Spirurida, *Rhabditida* (*Ancylostoma*), Diphyllobothriidea, Cyclophyllidea, *Cryptosporidium* and *Ascaririda*. We investigated the gut microbiome variation among samples by calculating the pairwise distances (β-diversity) based on the abundance (Bray-Curtis distances) of all identified cASVs. We used the package vegan v. 2.6-4 ^77^ with the function “distance”. Dissimilarity distances were then transposed to similarity distances (1-Bray-Curtis distances).

We tested the effect of individual repeatability on ß-diversity measures in 78 samples from 37 individuals in the overall microbiome and in restricted datasets of bacteria, fungi, and parasite members. We used distance-based intraclass correlation coefficients (dICC) and standard errors (SE) calculated based on 1000 bootstrap iterations, implemented with the package GUniFrac v.1.7 ^78^.

We tested for the association of immune measures while accounting for the effects of other known or expected host, social, and ecological variables on the β-diversity of the intestinal microbiome composition using dyadic comparisons (distances) among samples (excluding within-sample comparisons), as previously described ^39,79,80^. Host variables included age distance in days (Figure 1B), distances in f-IgA (Figure 1C), f-mucin levels (Figure 1D) and genetic mother. The ecological and social environment variables included distances in the standardised social rank, clan (same or different) and temporal distances (the distance in sample collection in days). We included an interaction between age and f-IgA and between age and f-mucin based on previous findings ^56^. We applied Bayesian generalised linear multilevel models using the Markov chain Monte Carlo algorithm No-U-Turn Sampler (NUTS)^81^ implemented in Stan through the brms package v. 2.19.0 ^82^. We used a multi-membership random-effects framework that accounts for individuals in each pairwise comparison (e.g., Individual A, Individual B). All predictors were scaled to values ranging from 0 to 1 to allow comparison of the standardised estimates of the predictors. We used four Markov chains, with 4 chains, 3000 iterations and 1000 burn-in iterations (warmup) to calibrate the Sampler, and default uninformative priors. We visually inspected convergence and assessed the relevance of each predictor by analysing R-hat values and the 95% credible intervals (95% CI). R-hat values provide information about chain convergence - when below 1.01, they were accepted as indicators of good convergence. A parameter was considered “significant” when the 95% CI did not include zero.

We applied a multivariate model to each microbiome component (parasites, fungi and bacteria), with the same predictors as the previous model and further captured the correlation between each group. We repeated all models using both occurrence i.e. presence/absence (Jaccard distances) and an alternative abundance-based ß-diversity measure (Aitchison distances), see Supplements, Table S1.

We applied a complementary method to further investigate the associations between immune measures (f-IgA and f-mucin) and overall microbiome composition. We implemented random forest regression, a tree-based ensemble machine learning in the package caret v. 6.0-94 ^83^, and implemented the ranger method in the ranger package v. 0.15.1 ^84^. The dataset was divided into training (80% of the samples) and test (the remaining 20%) sets using the function “createDataPartition”, that used stratified sampling to create the splits, ensuring a representative distribution of the respective immune measure between the sets. Training and tuning were performed with the function “trainControl” using 10-fold cross-validation which was repeated 10 times. The final values used for the model predicting f-IgA were mtry = 346, splitrule = variance min.node.size = 10, and for f-mucin were mtry = 188, splitrule = variance and min.node.size = 10. We then evaluated the predictions by using the resulting model to predict each immune measure in the test dataset and compare them with the observed values to calculate R^2^ from the respective regression model and Spearman correlation. Taxa importance was assessed based on permutation importance. We applied partial dependence plots using the package pdp v. 0.8.1 ^85^ to evaluate the marginal effects of the top 20 important taxa for the f-IgA and f-mucin predictions and visualise the direction and linearity of these associations (Supplements, Figures S2 and S3). Many taxa were not classified at phylum or even domain levels. Thus, we confirmed the taxonomy of these taxa for each immune measure by searching for the most similar nucleotide sequences using NCBI BLAST searches ^86^ and by inspecting the top hits (E-value, Table S3).

Co-abundance network analysis was used to infer potential ecologically relevant taxa associations. We filtered cASVs to 20% prevalence (143 cASVs remained) to reduce sparsity and ensure robustness. Co-abundance networks were created with SParse InversE Covariance estimation for Ecological Association Inference (SPIEC-EASI), with the package SpiecEasi ^87^ using the “mb” neighbourhood selection method. The optimal lambda coefficient for the network was 0.316. Network visualisation was done using the package igraph version 1.3.1 ^88^.

## Results

### Intestinal communities of free-ranging hyenas

We profiled the intestinal community of 158 free-ranging hyenas from three clans in the Serengeti National Park, sampled from 2004 to 2018. Thirty-seven individuals were sampled multiple times throughout their lives (Figure 1A). The median age of the animals at the time of sampling was 180 days, ranging from 52 days (1.7 months) to 5736 days (15.7 years). All adults were female (61 samples from 58 individuals) and juveniles were of both sexes (138 samples from 41 males and 81 females).

The overall intestinal community was composed of 999 cASVs deduced from 3133 ASVs across 35 different amplicons. 301 cASVs originated from bacteria and 691 cASV from eukarya (Figure 2A-D). Out of the latter, we identified 420 cASVs as fungi and 25 cASVs as known eukaryotic parasites of spotted hyenas annotated as Rhabditida (*Ancylostoma*) (6 cASVs), *Sarcocystis* (4 cASVs), Spirurida (3 cASVs), *Cystoisospora* (4 cASVs), *Cryptosporidium* (4 cASVs), Ascaridida (1 cASV), Diphyllobothriidea (1 cASV), Cyclophyllidea (2 cASVs). All but 5 samples had at least one parasite (cASV).

**Figure 2.**
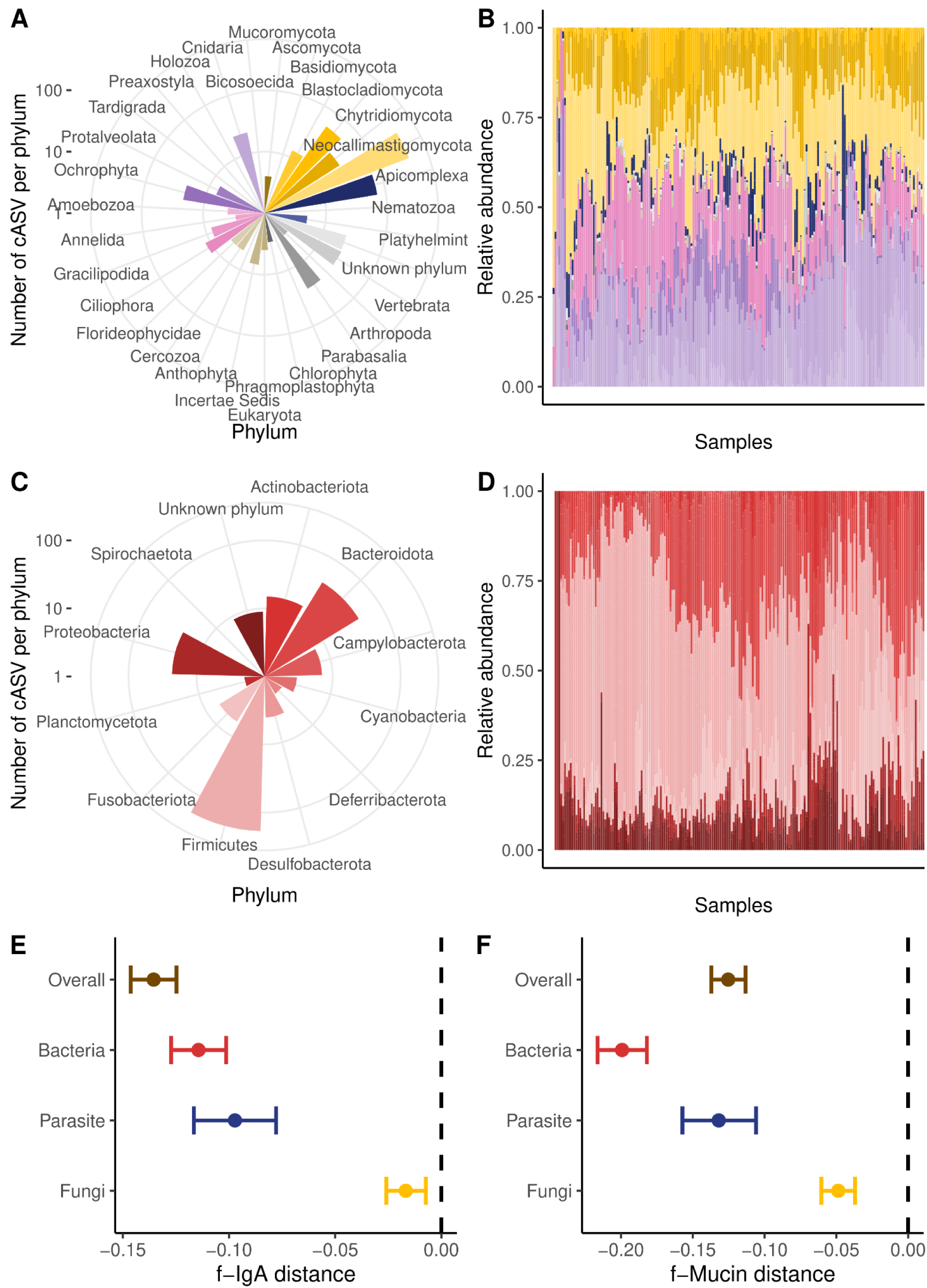
The intestinal microbiome composition of 199 samples from 158 spotted hyenas. **A)** and **B)** Eukaryote domain, **C)** and **D)** Bacterial domain. **A)** and **C)** depict the number of combined amplicon sequence variants (cASV) per phylum, **B)** and **D)** represent the relative abundance of each cASV (y-axis) for each sample (x-axis). Samples are sorted by clustering of Bray-Curtis dissimilarity. Colours represent the different phyla, with fungi in yellow, parasites in blue and bacteria in red. **E)** and **F)** Immune measures and age are strong predictors of intestinal microbiome composition. The compositional differences were analysed between samples using a Bayesian regression model with Bray-Curtis similarity estimates of ß-diversity as independent variables. The figure shows the posterior distributions of the predictor variables: **E)** faecal IgA (f-IgA) level distances; **F)** faecal mucin (f-mucin) level distances. The overall model consists of the overall compositional differences among whole intestinal communities (999 combined amplicon sequence variants (cASVs), brown). The decomposed communities were further analysed: bacteria (301 cASVs, red), fungi (420 cASVs, yellow), and parasites (25 cASVs, blue). Dots represent mean effect sizes, and estimates and bars represent corresponding 95% credible interval (CI) on intestinal community composition similarity. Note that the ranges on the x-axis differ between the plots, reflecting different effect sizes.

The repeatability within individuals tested multiple times was low or null for the overall intestinal microbiome (dICC = 0.084, SE = 0.037), bacteria (dICC = 0.061, SE = 0.049), fungi (dICC = -0.005, SE = 0.021), and highest for parasite composition (dICC = 0.108, SE = 0.080).

### Host immune measures are strongly associated with intestinal microbiome

We found a strong effect of the faecal immune measures on the overall intestinal microbiome (Table 1). While controlling for genetic mother, social rank, clan membership, and temporal effects, animal-pairs with similar f-IgA or f-mucin levels show significantly more similar compositions of the overall microbiome than animals with different mucosal immune levels (Table 1, Figure 2E,F).

**Table 1.** The effects of host-related predictors (immune measures, age) and social and ecological environments on the overall and bacteria, fungi, or parasite-specific intestinal community composition similarity (Bray-Curtis similarity distance) based on Bayesian regression multi-membership models. We show the mean estimates of the posterior distribution for each predictor and the associated 95% credible intervals (CI). All predictors are expressed as distances between compared pairs (n=19701). Effect sizes in bold indicate significant predictors for which both upper and lower 95% credible intervals (CI) are either positive or negative.

|  | Overall | Bacteria | Fungi | Parasite |
| --- | --- | --- | --- | --- |
| *Predictors* | *Estimate* | *Estimate* | *Estimate* | *Estimate* |
|  | *(credible interval)* | *(credible interval)* | *(credible interval)* | *(credible interval)* |
| f-IgA dist | **-0.060** | **-0.117** | **-0.013** | **-0.077** |
|  | **(-0.072 – -0.048)** | **(-0.135 – -0.099)** | **(-0.025 – -0.001)** | **(-0.103 – -0.052)** |
| f-mucin dist | **-0.125** | **-0.199** | **-0.049** | **-0.132** |
|  | **(-0.137 – -0.113)** | **(-0.216 – -0.182)** | **(-0.061 – -0.037)** | **(-0.157 – -0.106)** |
| Age dist | **-0.171** | **-0.195** | 0.004 | **-0.154** |
|  | **(-0.183 – -0.159)** | **(-0.212 – -0.177)** | (-0.009 – 0.016) | **(-0.180 – -0.127)** |
| Social rank | 0.002 | 0.001 | -0.001 | 0.001 |
| [same] | (-0.003 – 0.007) | (-0.006 – 0.008) | (-0.006 – 0.004) | (-0.009 – 0.012) |
| Genetic mother | 0.008 | 0.009 | 0.007 | **0.026** |
| [same] | (-0.003 – 0.019) | (-0.007 – 0.025) | (-0.004 – 0.018) | **(0.002 – 0.050)** |
| Temporal dist | **-0.012**<br>**(-0.017 – -0.006)** | **-0.012**<br>**(-0.020 – -0.004)** | **-0.032**<br>**(-0.038 – -0.027)** | **-0.018**<br>**(-0.029 – -0.006)** |
| Clan [same] | 0.002<br>(-0.000 – 0.005) | 0.003<br>(-0.001 – 0.006) | -0.000<br>(-0.003 – 0.002) | -0.003<br>(-0.008 – 0.002) |
| f-IgA dist: Age | **0.064** | -0.020 | **-0.046** | **0.115** |
| dist | **(0.031 – 0.097)** | (-0.068 – 0.027) | **(-0.078 – -0.014)** | **(0.045 – 0.184)** |
| f-mucin dist: Age | **0.090** | -0.016 | **0.050** | **0.146** |
| dist | **(0.059 – 0.121)** | (-0.061 – 0.030) | **(0.019 – 0.080)** | **(0.082 – 0.213)** |

Age was the most important predictor of overall microbiome composition (Table 1). The similarity of the overall microbiome and the compositions of bacteria and parasites decreased with increasing age distances (Table 1, Figure 3A). The effects of immune measures depended on age for the overall microbiome, parasite, and fungi compositions (Table 1, Figure 3B-C). When pairs of samples from individuals of similar age were compared (purple lines in Figure 3D-E), the microbiome similarities (overall and parasite compositions) decreased with increasing f-IgA and f-mucin distances. When pairs with large age differences were compared (green-blue lines in Figure 3D-E), this effect became slightly less pronounced than in pairs with small age differences. On the contrary, fungi composition similarity became more pronounced with increasing f-IgA distances, but not f-mucin distances, than in pairs with small age differences (Figure 3B-C).

**Figure 3.**
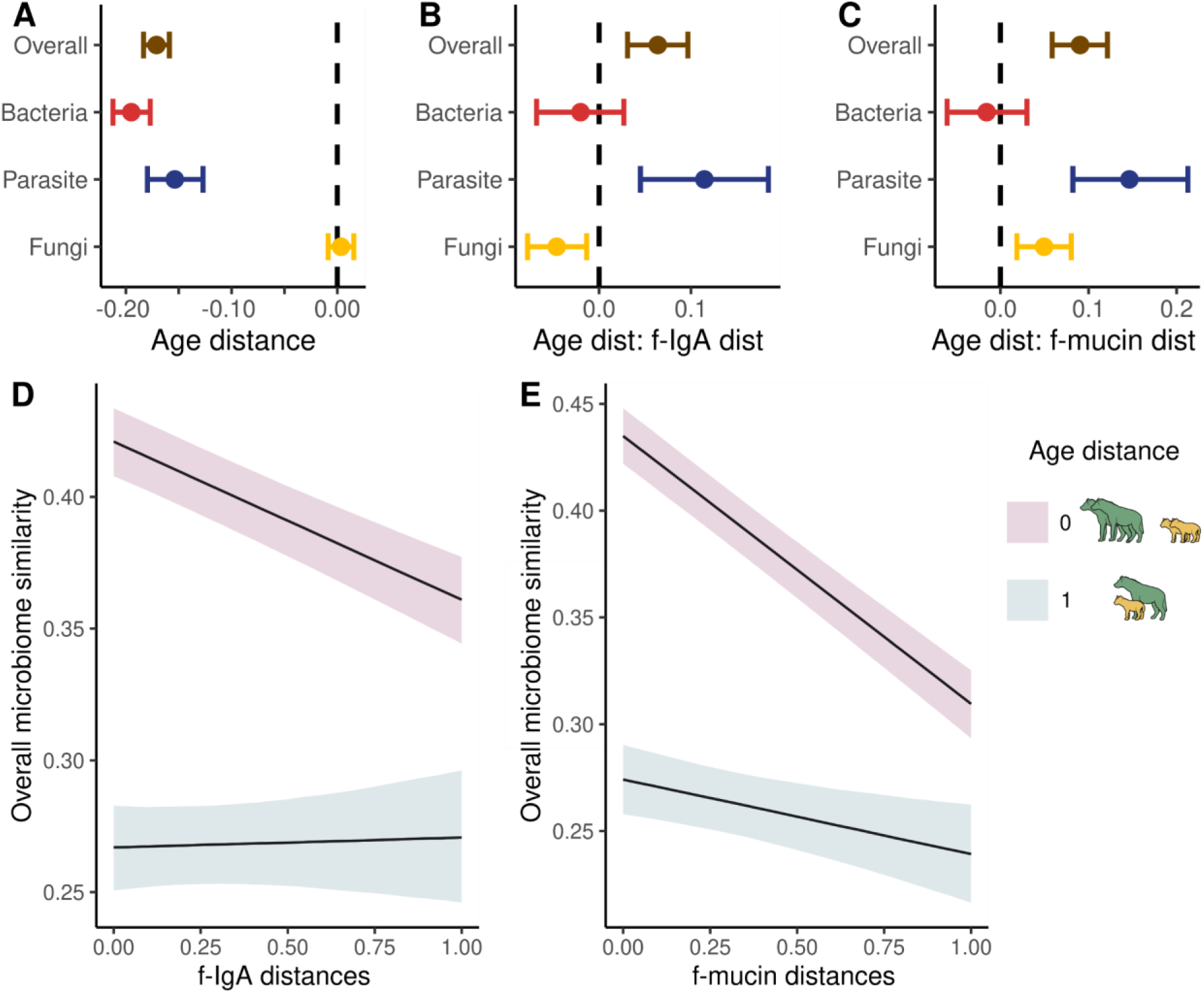
Similarity of the overall intestinal community composition is mediated by age. **A-C**) The compositional differences were analysed between samples using a Bayesian regression model with Bray-Curtis similarity estimates of ß-diversity as independent variables. Age distances are important for all but the fungal members of the intestinal community. Shown are the posterior distributions of **A**) age, **B**) age interacting with f-IgA, and **C**) age interacting with f-mucin distances. The predicted similarity of the intestinal microbiome is displayed between sample pairs with similar animal age (age distance = 0) and in pairs with extreme age distances (age distance = 1) varying with **D**) f-IgA and **D**) f-mucin distances. The black lines indicate the mean predicted immune measures and the shaded area represent the 95% credible intervals (CI).

Temporal distances between the time of sampling, i.e. a large amount of time elapsed between the collection of two samples, decreased the microbiome similarity of the overall microbiome composition and all microbiome members (Table 1). Sharing the same genetic mother had a small but significant positive effect on parasite composition similarity.

### Host immune measures are predicted by the overall intestinal microbiome composition

To further investigate the association between the intestinal microbiome and mucosal immunity, we applied a random forest regression to the relative abundances of all 999 cASV identified to predict f-IgA and f-mucin. The 199 samples were divided into training (80%) and testing (20%) sets. Predicted and observed values correlated moderately for f-IgA (R^2^ = 0.352, Spearman’s rho = 0.599, p<0.001, n = 39, Figure 4A) and f-mucin (R^2^ = 0.311; Spearman’s rho = 0.556, p<0.001, n = 39, Figure 4C). We identified the highest ranking cASVs (top 20 cASVs) according to the random forest importance score for predicting f-IgA (Figure 4B) and f-mucin levels (Figure 4D). Sequences are available in additional file 2. Partial dependence plots were used to visualise the marginal effects for each of the highest-ranking cASVs for predicting f-IgA (Figure S2) and f-mucin levels (Figure S3).

**Figure 4.**
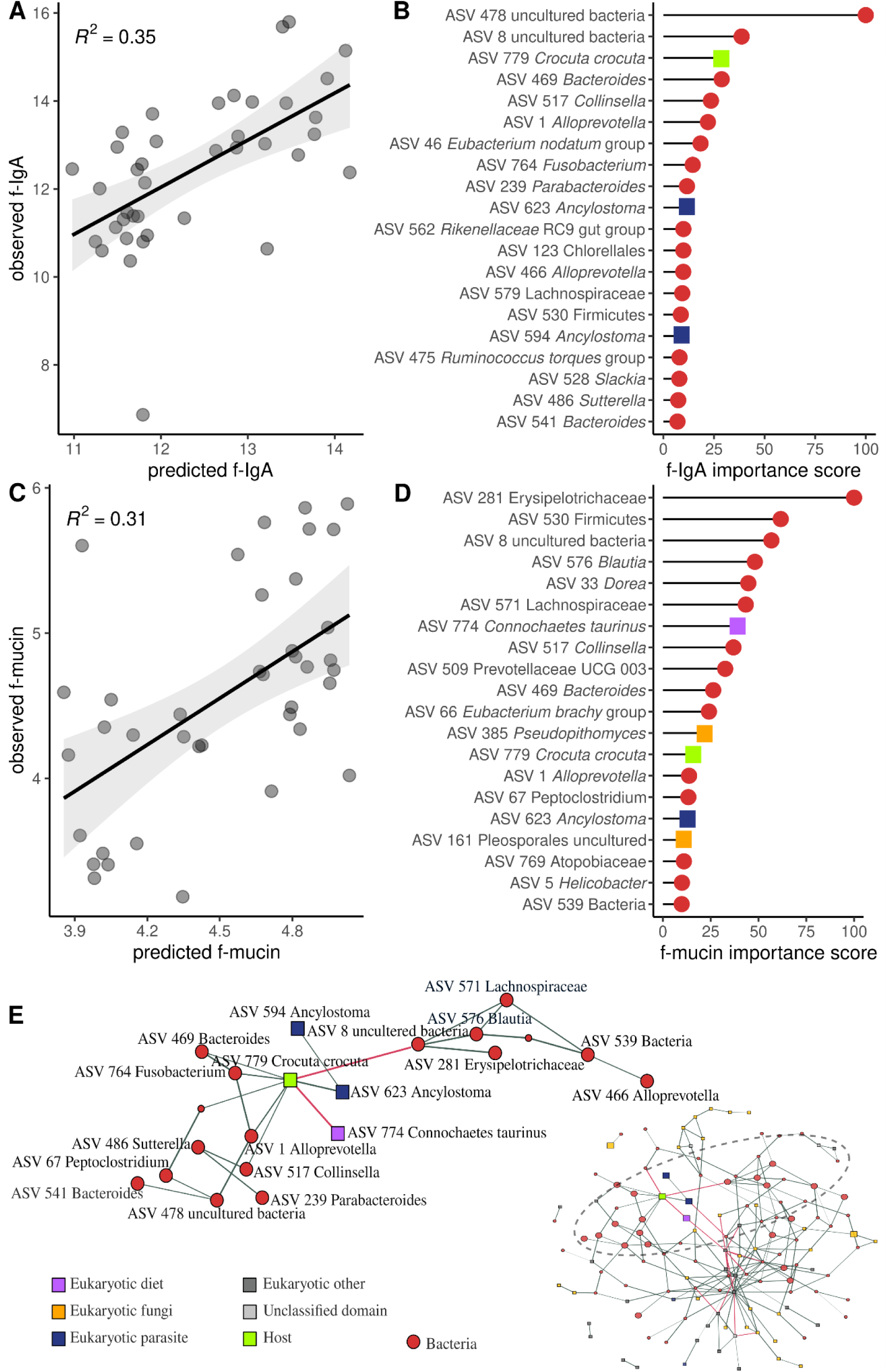
Overall microbiome abundance predicts faecal IgA (f-IgA) and mucin (f-mucin) levels in hyenas. Random forest regressions were implemented with 999 combined amplicon sequence variants (cASVs) from 160 faecal samples (training set) and evaluated with an independent dataset of 39 samples (test set), to predict **A**) and **B**) f-IgA levels and **C**) and **D**) f-mucin levels. The model accuracy was based on linear regression of predicted against observed **A**) f-IgA and **C**) f-mucin. Grey lines represent the regression lines from linear models calculated from the predicted against the observed respective immune measures, with the corresponding R^2^. The top 20 most important cASVs for model prediction of **B**) f-IgA and **D**) f-mucin levels. Dots are colour coloured. Red represents taxa from the bacterial domain, blue parasites, yellow fungi, green host DNA, and lilac prey (diet) DNA. **E)** Co-occurrence network of 143 taxa across 199 samples from 158 spotted hyenas. Nodes represent a cASV and edges the association between a pair of cASVs. The edge thickness is scaled by association strength. The green and red edges represent positive and negative associations, respectively. The most important taxa for predicting both f-IgA and f-mucin are depicted in a larger node size. The area under the dashed ellipse is highlighted.

Among the highest-ranking predictors of f-IgA levels were 17 bacteria cASV, host DNA (cASV 776 *Crocuta crocuta)* and 2 parasite cASVs (both *Ancylostoma*). For f-mucin levels, the highest-ranking predictors were 15 bacteria cASVs, host DNA (cASV 779 *Crocuta crocuta),* prey DNA (*cASV* 774 *Chonnochaetes taurinus)*, one parasite cASV (cASV 623 *Ancylostoma*) and two fungi cASV (cASV 385 *Pseudopithomyces* and cASV 161 Pleosporales uncultured). Out of these, the same 7 cASV were predicting both f-IgA and f- mucin levels: cASV 8 (uncultured bacteria), cASV 779 *Crocuta crocuta*, cASV *469 Bacteroides*, cASV 517 *Collinsella*, cASV 1 *Alloprevotella*, and cASV 623 *Ancylostoma*.

### Inter-taxa associations

Among compared animals, fungi composition similarity increased with parasite similarity (residual correlation estimate (CI) = 0.07 (0.06 - 0.09), and with bacteria similarity (residual correlation estimate = 0.12 (0.11 - 0.13) and bacteria similarity increased with parasite similarity (residual correlation estimate = 0.17 (0.16 - 0.19). We further investigated inter-taxa interactions with a co-abundance network of cASVs present in at least 20% of the samples. The resulting network contained 143 nodes and 219 edges (Figure 4E). The resulting network reflects a highly connected community, with the most important taxa for predicting both f-IgA and f-mucin being closely associated.

## Discussion

The mammalian mucosal immunity and gut microbiome are in constant cross-talk. The resulting interactions are influenced by host characteristics and by the host’s biotic and abiotic environments. Knowledge on symbiont-symbiont and symbiont-immune interactions is limited, especially outside the scope of human and model organism studies. Wild populations are exposed to and harbour a much more diverse array of macro- and microorganisms, live in heterogeneous environments and have diverse genetic backgrounds. Here, we explore the associations between two important and broad-acting measures of intestinal mucosal immunity, IgA and mucin, and the gut microbiome, while accounting for host characteristics, social rank, and environmental factors in Serengeti hyenas. Both IgA and mucin were strongly associated with the intestinal community and these associations varied in strength within different components of the gut microbiome, being stronger within bacteria, intermediate within parasites and weaker within fungi communities. The most important taxa predicting both immune measures were predominately bacteria and the parasite *Ancylostoma.* Our results link, for the first to our knowledge, mucosal immunity to intestinal microbiome composition, considering both bacteria and eukaryotes, in a wild population.

Bacteria had stronger associations with both IgA and mucin than parasites and fungi, regardless of host characteristics and the environment. Through a complementary analysis, we confirmed that the composition of the gut microbiome, particularly bacteria, predicted the levels of f-IgA and f-mucin well. This is not surprising, as bacteria outnumber eukaryotes in mammalian intestines ^89,28,90^ and in the metabolic products that are currently known to interact with the immune system ^91,92^. Parasites were disproportionately associated with both immune measures, considering that they comprise fewer taxa than bacteria and fungi. This is expected, because parasites can tamper with immune responses to their own advantage, e.g. by secreting enzymes that degrade the mucin layer ^93^. The association between fungi and mucosal immunity is still poorly understood, but recent studies have revealed potentially relevant associations ^22,94^.

Many of the identified predictive taxa are well-known intestinal symbionts in humans and mouse models, associated with either health benefits or detriments. Some of these bacterial taxa, including *Bacteroides, Alloprevotella, Peptoclostridium*, *Slackia,* and *Fusobacterium*, were previously identified as ‘core gut symbionts’, i.e. present in 85% of samples from a different population of spotted hyenas ^55^. It is conceivable that these taxa provide relevant functions to hyenas. The *Bacteroides* genus is a common symbiont of humans ^95^ and other mammals ^96–98^ and its members have been associated with beneficial ^99^ and negative effects in their host ^100^. Although the functional role in intestinal health is still largely unknown, *Alloprevotella* is suggested to have anti-inflammatory properties in humans ^101^, is reduced after a stressful intervention in pigs (*Sus domesticus*) ^102^, and is associated with the presence of helminths in wild chimpanzees (*Pan troglodytes*) ^103^. Interestingly, two taxa from *Alloprevotella* abundance positively and negatively predicted f-IgA levels in our study, which might point to species or possibly strain-specific regulation by IgA in this genus. Taxa of the genus *Slackia* may exert an indirect anti-inflammatory effect in the gut via metabolite production that can maintain immunity homeostasis ^104^. Taxa of the *Fusobacterium* genus are associated with disease states in humans and in its translational models. *F. nucleatum* causes inflammation by upregulating the activity of IgA in laboratory mice ^105,106^ and worsens colorectal cancer in humans ^107,108^. Members of the genus *Peptoclostridium* can help maintain the intestinal barrier, among other functions ^109^, but can also be life-threatening to humans and other animals ^110^.

We found the parasite *Ancylostoma* to be positively associated with both f-IgA and f-mucin levels. *Ancylostoma* is a blood-feeding nematode that causes extensive inflammation and intestinal bleeding in several different species ^111–113^. Our findings are consistent with those of a previous study in the same population, in which *Ancylostoma* egg load was found to be positively associated with both f-mucin and f-IgA ^56^. We interpret this result as the potential upregulation of IgA and mucin by the host in an unsuccessful attempt to eliminate adult *Ancylostoma* parasites from the gut. The presence of host DNA is also positively associated with f-IgA and f-mucin, which could indicate intestinal inflammation because of increased cell shedding leading to higher levels of host DNA in faeces ^114,115^. In the co-occurrence network, host DNA is positively associated with both *Ancylostoma* and *Fusobacterium,* both of which are capable of causing extensive inflammation ^111–113,108^. It is also possible that concentrations of IgA, mucin, and host DNA co-varied with intestinal transit or diet ^116,117^ and could be indicators of the host’s normal variation through physiological states, although our results indicate that prey DNA is negatively correlated with host DNA, and host DNA correlated with a known parasite in hyenas, thus rather pointing to disruption or disease.

Our co-abundance network and inter-taxa correlations suggest a highly connected intestinal community. The taxa predicting f-IgA and f-mucin are closely associated, which potentially indicates a regulatory role of the mucosal immune system. Previous studies in wild non-human primates have also documented associations between parasites and bacteria, which are shaped by host-related and ecological factors ^118–120^. The role and significance of fungi within the intestinal community remains to be elucidated given the scarcity of studies focusing on this taxonomic group within wildlife guts, but see ^35,118^.

As expected from previous studies on hyenas ^30,56^ and other wild animals (e.g. wild primates ^121^), we found that age is a strong predictor of microbiome similarity. Our results suggest that the strength of the associations between bacteria, parasites, and fungi on the one hand and mucosal immune measures on the other were modulated by host age. These effects were complex and inconsistent between measures of mucosal immunity and among the different components of the gut microbiome. The mammalian gastrointestinal tract is colonised at birth by pioneer microbes acquired from mothers and the environment ^122,123^ and the gut microbiome undergoes a process of microbial succession ^124^. This process is closely linked to the maturation of the immune system, as the immune system requires microbial interactions early in life for proper development and maturation, and in turn shapes the microbiome composition ^8,125^. Our results are consistent with complex and context-specific interactions between the immune response and the host throughout life.

It is difficult to comment on the health effects of a particular microbiome community or even on those of individual taxon and associations, other than known parasites such as *Ancylostoma* ^56^. Thus, we are cautious when interpreting the functional roles of individual taxa associated with mucosal immunity in this study. Different species, strains, and immunogenic potential within these taxa could lead to different outcomes in wild populations for which knowledge is still limited. Host-microbial interactions are context-dependent and are mostly studied in humans and laboratory translational models. In future studies, assessments of the effects of particular microbial communities on individual health (as measured using e.g. body condition) may help determine whether specific taxa are beneficial or not to hosts, and thus involved in shaping host evolutionary fitness. Because we measured broad-acting measures of mucosal immunity, we cannot infer specific mechanisms of interactions between the different members of the gut microbiome and the host. In the future, measuring antigen-specific IgA isoforms and mucin types together with assessments of intestinal communities may shed light on the finer details of the regulation of individual taxa in the biomes. In the broad absence of reagents for wildlife antibody detection ^126^, IgA-seq, the enrichment of sequencing of IgA-coated taxa ^127,128^ could provide unprecedented insights into taxon-specific immune control at intestinal barriers. Investigating the link between host fitness, the microbiome and its metabolites could reveal evolutionary adaptations that promote or hamper host health and performance.

Natural populations harbour a hidden and mostly unknown diversity within their guts, and their immune systems must regulate such communities; maintaining mutualistic and commensals and reducing detrimental parasitic interactions. We identified broad and general associations between immune measures and the different members of the microbiome and pinpointed the taxa driving these associations. These findings indicate the important role that the immune system plays in the defence and also regulation of the microbiome, and we propose that the identified taxa are closely associated with and involved in the cross-talk within the gut of natural populations of hyenas - a potential product of co-adaptation. The next step is to further investigate the genetic diversity and functional profiling of gut microbiomes in natural populations to uncover evolutionary aspects of such potential co-adaptations. We thus encourage others to approach wildlife microbiome research in a holistic manner and incorporate the measures of the immune system, both systemically and at mucosal sites, to improve our understanding of the complex and dynamic host-microbiome interactions.

## Author contribution

EH, GAC, SB, AW, MLE and HH conceptualised the original study and acquired funding. SCMF, SPVS, EH and SB designed the analyses and computational framework. SCMF, SM, MLE, HH and SB conducted fieldwork. SCMF and MV performed the laboratory work for immune measures. VHJD and SK performed laboratory work for microbiome analyses. SCMF and EH analysed the data. SCMF and SPVS wrote the manuscript with contribution and feedback from SB and EH. All authors contributed significantly to editing the manuscript.

## Availability of data and materials

The datasets supporting the conclusions of this article are available in the GitHub repository: https://github.com/ferreira-scm/Microbiome_Hyena.git. All sequencing raw data can be accessed through the BioProject PRJNA1134446 in the NCBI SRA.

## Acknowledgments

For fieldwork, we were granted research permits from the Tanzania Commission for Science and Technology (COSTECH) and permission from the Tanzanian National Parks Authority (TANAPA) and Tanzanian Wildlife Research Institute (TAWIRI). Fieldwork was supported by the Commission for Science and Technology of Tanzania (COSTECH), the Tanzania Wildlife Research Institute (TAWIRI), and Tanzania National Parks (TANAPA). For laboratory support at the Leibniz Institute for Zoo and Wildlife Research (IZW), we thank D. Thierer, K. Pohle, F. Webster and C. Bost. We thank A. Francis, T. Shabani, M. Andris, N. Boyer, T. Golla, K. Goller, N. Gusset-Burgener, B. Kostka, M. Lindson, D. Thierer, A. Türk and K. Wilhelm for field and technical assistance.

All procedures and protocols were developed and implemented in compliance with the Leibniz Institute for Zoo and Wildlife Research Ethics Committee on Animal Welfare (permit number: 2017-11-02). Hyena icons were designed by Sonja Metzger.

