## Supplementary information for "Spotted hyena gut cross-talks: Symbionts modulate mucosal immunity"

* corresponding author(s)

### Supplementary methods

#### Faecal immunological assays

The faecal mucin assay detects and quantifies oligosaccharides released from mucin by discriminating between O-linked and N-linked glycoproteins (Crowther & Wetmore, 1987; Bovee-Oudenhoven et al., 1996) in freeze-dried samples. Briefly, freeze-dried samples were suspended in phosphate-buffered saline (PBS) solution and, then mixed and incubated in a shaking bath to denature glycosidases. Incubation was followed by centrifugation for solubilisation and separation. The resulting supernatant was mixed with alkaline 2-cyanoacetamide (CNA), and incubated. Borate buffer was added, and the solution was cooled to room temperature. The measurements were conducted using a fluorometric assay at 383 nm with an excitatory wavelength of 336 nm (Infinite M200, TECAN, Männedorf, Switzerland), with a low-binding 96-well plate (PerkinElmer(R)). Results are expressed as µmol oligosaccharide equivalents to a standard curve created with N-acetylgalactosamine (Sigma-Aldrich(R), Darnstadt, Germany), whereas porcine stomach mucin was used as a positive control (Sigma-Aldrich(R), Darmstadt, Germany).

f-IgA levels were measured using a sandwich ELISA altered from (Tress et al., 2006). Briefly, saline extracts from freeze-dried faecal samples were used (Ferguson et al., 1995; Peters et al., 2004), and protease-inhibitor MixM (Serva Electrophoresis GmbH, Heidelberg, Germany) was added. The resulting supernatant was stored at -20°C until use. For the sandwich ELISA, an anti-cat IgA (Lot.A10, Novusbio, Abingdon, UK) was used as the capture antibody, and conjugated anti-cat IgA (Lot.P18, Novusbio, Abingdon, UK) as detection antibody. The plates were read at a wavelength of 450 nm using the BioTek Quant Microplate reader (BioTek, Vermont, USA). The results are presented as relative units (RU), and standard curves were obtained with a pool of 72 samples. For both mucin and f-IgA assays, all plates included negative controls and quality controls, and all samples were performed in duplicate and results were accepted if the coefficient of variation was below 5% and within the working range previously established (Ferreira et al., 2021).

### Supplementary figures

**Figure S1**. Sampling design of the study. a) Frequency count of the number of sampled siblings per individual included in this study. b) Frequency count of sampled individuals belonging to each clan (I, M and P). c) Histogram of the individual standardised social rank across samples. d) Histogram of the year of sampling across samples. Individual count includes 158 individuals and sample count includes 199 samples.

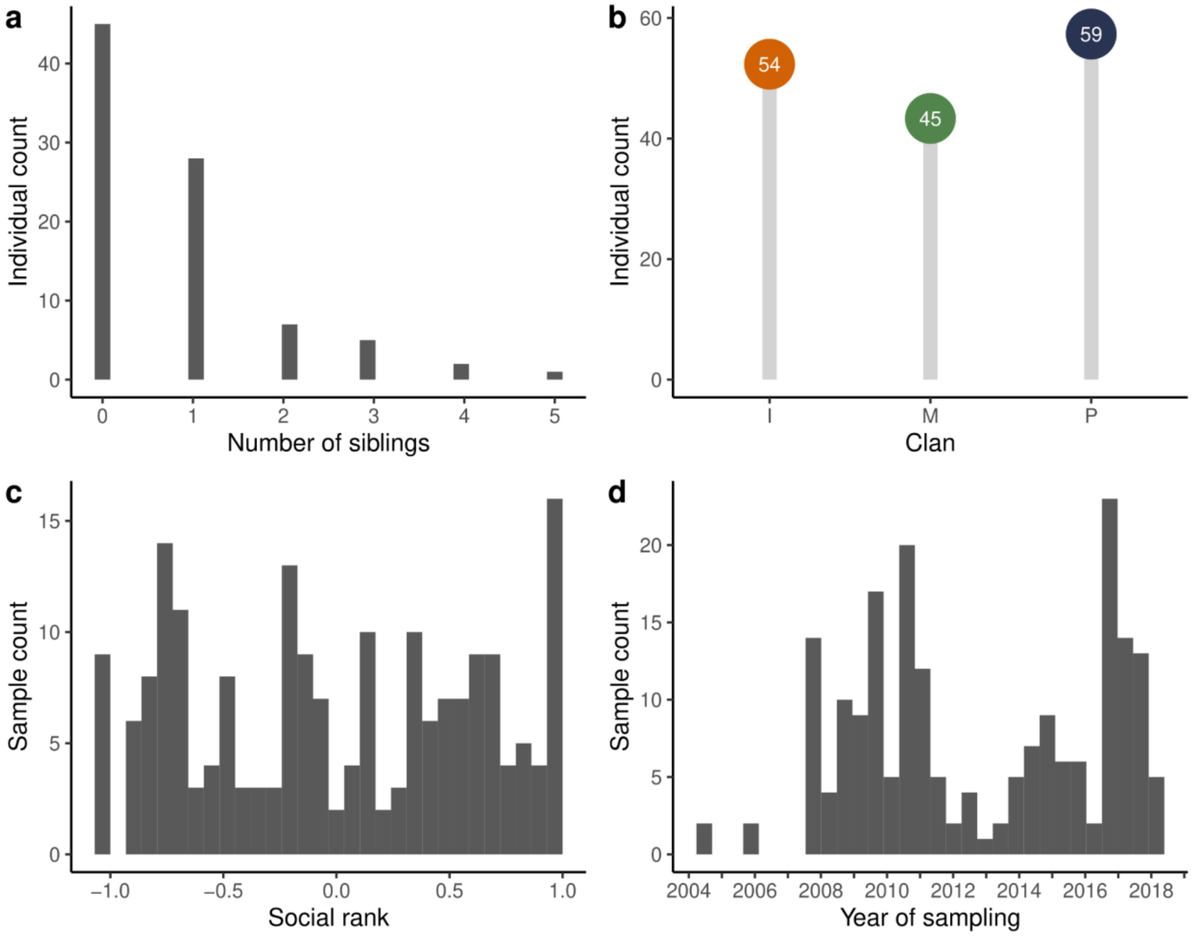

**Fig.S2 Marginal effect size of the 20 most important taxa for prediction of f-IgA.** The relationship between f-IgA and the top 20 cASVs based on partial dependence plots. On the y-axis, yhat is expected f-IgA as a function of the relative abundances of cASVs. Red represents taxa from the bacterial domain; brown, the host DNA and blue, eukaryotic parasite taxa.

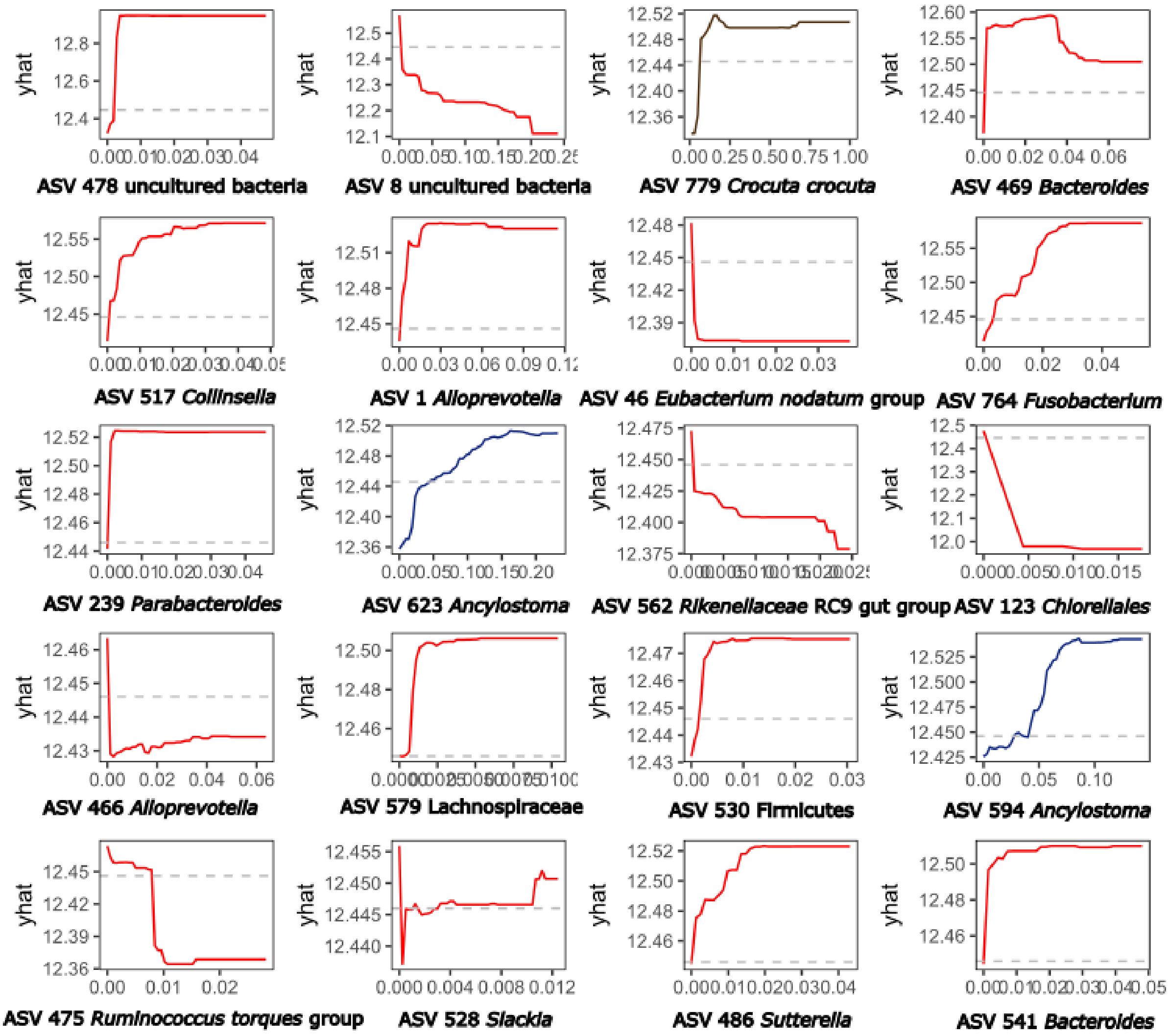

**Fig.S3 Marginal effect size of the 20 most important taxa for prediction of f-mucin.** The relationship between f-mucin and the top 20 cASVs based on partial dependence plots. On the y-axis yhat is expected f-mucin as a function of the relative abundances of cASVs. Red represents taxa from the bacterial domain; brown, the host DNA; blue, eukaryotic parasite taxa; lilac, prey DNA; and yellow, fungi.

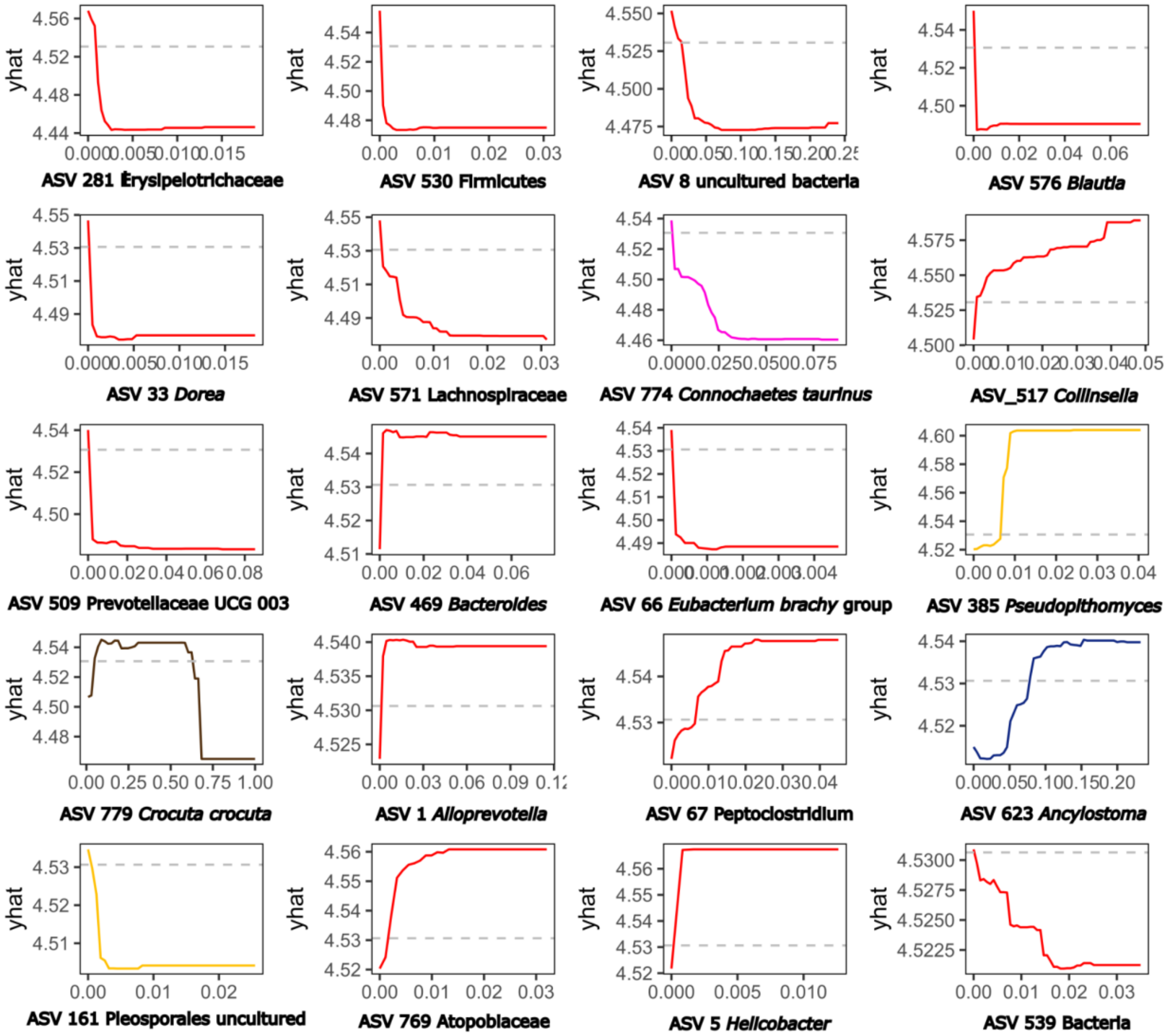

**Table S1.** Host and ecological factors affect the intestinal community composition similarity (Jaccard and Aitchison similarity distance). Shown are the Bayesian regression multi-membership models testing the effect of each predictor on microbiome similarities. We show the mean estimates of the posterior distribution for each predictor and the associated 95% credible intervals (95% CI). R-hat values provide information on the chain convergence and are all below 1.01, indicating good convergence. A parameter is considered significant when the 95% CI does not include zero and is shown as coloured. All predictors are expressed in distances between compared pairs, n=19701. Brown represents the effect size on the overall community; blue, on parasites; yellow, on fungi and red, on bacteria members of the microbiome.

| Jaccard similarity distances (occurrence-based) | | | | | | | | | | |
|---|---|---|---|---|---|---|---|---|---|---|
| | | Age dist | f-IgA dist | f-Mucin dist | Social rank dist | Genetic mother [same] | Temporal dist | Clan [same] | Age:f-IgA | Age:f-mucin |
| Overall | Estimate | -0.106 | -0.056 | -0.107 | 0.001 | 0.003 | -0.011 | 0.002 | -0.027 | 0.001 |
| | 95% CI | -0.115: -0.097 | -0.065: -0.047 | -0.116: -0.098 | -0.003: 0.004 | -0.006: 0.011 | -0.015: -0.007 | 0.000: 0.004 | -0.051: -0.003 | -0.021: 0.024 |
| Parasite | Estimate | -0.233 | -0.068 | -0.031 | -0.003 | 0.012 | -0.016 | 0.002 | 0.184 | 0.022 |
| | 95% CI | -0.255: -0.210 | -0.091: -0.046 | -0.055: -0.008 | -0.013: 0.006 | -0.009: 0.033 | -0.027: -0.006 | -0.003: 0.007 | 0.122: 0.246 | -0.036: 0.081 |
| Fungi | Estimate | -0.001 | -0.013 | -0.040 | -0.002 | 0.006 | -0.014 | 0.001 | -0.022 | 0.033 |
| | 95% CI | -0.010: 0.007 | -0.022: -0.005 | -0.048: -0.031 | -0.005: 0.001 | -0.002: 0.014 | -0.018: -0.010 | -0.001: 0.003 | -0.044: 0.001 | 0.011: 0.054 |
| Bacteria | Estimate | -0.197 | -0.108 | -0.192 | 0.001 | -0.001 | -0.007 | 0.001 | -0.061 | -0.028 |
| | 95% CI | -0.213: -0.181 | -0.124: -0.092 | -0.208: -0.176 | -0.005: 0.008 | -0.015: 0.015 | -0.015: 0.000 | -0.002: 0.005 | -0.105: -0.018 | -0.070: 0.013 |
| Aitchison similarity distances (abundance-based) | | | | | | | | | | |
| | | Age dist | f-IgA dist | f-Mucin dist | Social rank dist | Genetic mother [same] | Temporal dist | Clan [same] | Age:f-IgA | Age:f-mucin |
| Overall | Estimate | -0.136 | -0.057 | -0.067 | 0.002 | 0.007 | -0.012 | 0.002 | 0.039 | 0.034 |
| | 95% CI | -0.146: -0.125 | -0.068: -0.046 | -0.078: -0.056 | -0.002: 0.007 | -0.003: 0.017 | -0.017: -0.007 | -0.000: 0.004 | 0.011: 0.069 | 0.006: 0.063 |
| Parasite | Estimate | -0.097 | -0.100 | -0.047 | 0.005 | 0.027 | -0.012 | -0.003 | 0.072 | 0.066 |
| | 95% CI | -0.116: -0.077 | -0.120: -0.080 | -0.066: -0.028 | -0.003: 0.012 | 0.009: 0.044 | -0.020: -0.003 | -0.006: 0.001 | 0.019: 0.124 | 0.017: 0.114 |
| Fungi | Estimate | -0.019 | 0.020 | -0.009 | 0.002 | 0.004 | -0.021 | -0.001 | 0.055 | 0.017 |

| | | | | | | | | | | |
|---|---|---|---|---|---|---|---|---|---|---|
| | 95% CI | -0.028: -0.010 | 0.010: 0.030 | -0.019: -0.000 | -0.001: 0.006 | -0.005: 0.012 | -0.025: -0.016 | -0.002: 0.001 | 0.030: 0.081 | -0.007: 0.041 |
| Bacteria | Estimate | -0.116 | -0.075 | -0.094 | 0.000 | 0.010 | -0.005 | 0.002 | 0.016 | 0.002 |
| | 95% CI | -0.129: -0.102 | -0.090: -0.061 | -0.107: -0.080 | -0.005: 0.006 | -0.002: 0.022 | -0.011: 0.001 | -0.000: 0.005 | -0.021: 0.054 | -0.034: 0.036 |
